# Synthetic Coolant WS-23 increases E-Cigarette Generated Aerosolized Acellular Reactive Oxygen Species (ROS) Levels

**DOI:** 10.1101/2022.06.20.496866

**Authors:** Shaiesh Yogeswaran, Marko Manevski, Hitendra S. Chand, Irfan Rahman

## Abstract

There has been a substantial rise in e-cigarette (e-cig) use or vaping in the past decade, prompting growing concerns about their adverse health effects. Recently, e-cig manufacturers have been using synthetic cooling agents, like WS-23 and WS-3, to provide a cooling sensation without the “menthol taste”. Studies have shown that aerosols/vapes generated by e-cigs can contain significant levels of reactive oxygen species (ROS). However, studies investigating the role of synthetic coolants in modulating ROS levels generated by e-cigs are lacking. This study seeks to understand the potential of synthetic coolants, e-cigarette additives that have become increasingly prevalent in e-liquids sold in the United States (US), on acellular ROS production. Aerosols were generated from e-liquids with and without synthetic coolants through a single-puff aerosol generator; subsequently, acellular ROS was semi-quantified in H2O2 equivalents via fluorescence spectroscopy. Our data suggest that adding WS-3 to e-liquid base (PG:VG), regardless of nicotine content, has a minimal impact on modifying e-cigarette-generated acellular ROS levels. Additionally, our data also suggest that the addition of WS-23 to nicotine-containing e-liquid base significantly modifies e-cigarette-generated acellular ROS levels. Together, our data provide insight into whether adding synthetic coolants to e-liquids significantly impacts vaping-induced oxidative stress in the lungs.

## 1. Introduction

During the past few years, adolescent use of e-cigs or various electronic nicotine delivery systems (ENDS) has significantly increased, thus leading to an increase in the prevalence of E-cigarette or Vaping Associated Lung injury (EVALI) across the United States (King, Jones et al. 2020). As of February 18, 2020, a total of 2,807 EVALI-related hospitalizations or deaths were reported to the Centers for Disease Control (CDC) from all 50 states (King, Jones et al. 2020). Consequently, the Food & Drug Administration (FDA) implemented an e-cigarette flavor enforcement policy banning the sales of all flavored cartridge-based nicotine-containing e-cigarette products, excluding tobacco and menthol flavors (Lu, Sun et al. 2022).

Following the FDA’s 2020 flavor-enforcement policy, menthol-flavored e-cigarette sales had significantly increased in the US; specifically, there was a 54.5% increase in the market share of menthol-flavored e-cigarettes over four weeks and an 82.8% increase over eight weeks following the FDA’s ruling (Diaz, Donovan et al. 2021). The cooling sensation created by menthol plays a significant role in the decision of both youth and adults to continue to vape, as it masks the bitter taste of nicotine (Davis, Morean et al. 2021). However, recently, more e-cigarette manufacturers have switched to non-menthol-containing flavoring chemicals to make e-cigarettes that give users a cooling sensation upon inhalation. These flavoring chemicals include synthetic coolants, like Methyl diisopropyl propionamide (WS-23) and N-Ethyl-2-isopropyl-5 methylcyclohexanecarboxamide (WS-3) (Davis, Morean et al. 2021, Jabba, Erythropel et al. 2022).

Examples of e-cigarette flavors containing WS-23 or WS-3 include e-cigarette flavors with “ice”, “chilled”, “cooled”, and “polar” in their name; some of these e-cigarette flavors consist of flavor combinations with fruity and drink flavors, like “melon-ice”, “blueberry-ice”, and “iced-pink punch” (Leventhal, Dai et al. 2021). The significant increase in the marketing of “iced/cooled” flavored e-cigarettes in the U.S had occurred right around the time when sales of disposable e-cigarettes surged following the FDA’s implementation of its March 2020 e-cigarette flavor enforcement policy (Leventhal, Dai et al. 2021). One lab found WS-23 to be a major component within the nicotine-containing e-liquid-pods, a type of ENDS, given to them by recovered EVALI patients in New York State (Lu, Li et al. 2021). Additionally, one study (Jabba, Erythropel et al. 2022) found that WS-23 was present in e-cigarettes marketed in the US at levels that may potentially result in exceeding the Margin of Exposure (MOE), a risk assessment parameter for toxic compounds used by World Health Organization (WHO) (Jabba, Erythropel et al. 2022). Jabba, Erythropel et al. 2022’s results suggest that those who use e-liquids comprised of W-3 or WS-23 are potentially at risk for long-term pulmonary health issues (Jabba, Erythropel et al. 2022).

Aerosols generated by e-cigarettes or other ENDS modalities have been found to contain dangerous chemicals, including formaldehyde and acetaldehyde, which are known to cause lung cancer and cardiovascular disease (Ogunwale, Li et al. 2017). Also, consistently, it has been found that dysregulated inflammatory cytokine output is an effect of chronic e-cig exposure in both *in vivo* and *in vitro* models (Davis, Sapey et al. 2022). Moreover, previous studies have shown that aerosols generated by flavored e-cigs produce significant levels of acellular reactive oxygen species (ROS) and induce cellular ROS in small airway epithelial cells (SAEC) (Zhao, Zhang et al. 2018, Yogeswaran, Muthumalage et al. 2021, Yogeswaran and Rahman 2022). ROS, either exogenous or when produced in excess endogenously, can lead to a redox imbalance in the lungs (Zuo and Wijegunawardana 2021). One study found tobacco smoke to contain a significant amount of free radicals, ∼1 × 10^15^ radicals per puff (Pryor and Stone 1993, Valavanidis, Vlachogianni et al. 2009, van der Toorn, Rezayat et al. 2009). ROS in smoke generated from conventional cigarettes, when inhaled, will react with antioxidants in the epithelial lining fluid (ELF) covering airway epithelial cells (Valavanidis, Vlachogianni et al. 2009). Moreover, ROS in tobacco smoke, after reaching the ELF of airways, can lead to the destruction of endogenous antioxidants, thus significantly reducing cellular antioxidant capacity (van der Toorn, Rezayat et al. 2009). Oxidative stress induced by this redox imbalance has been implicated in the pathology of many types of lung diseases, such as acute respiratory distress syndrome (ARDS), asthma, and chronic obstructive pulmonary disease (COPD) (Zuo and Wijegunawardana 2021).

Studies so far have shown that exposure to e-cigarette aerosols induces oxidative stress in the lungs (Wang, Zhang et al. 2020). Regarding ROS-related e-cigarette studies, studies have shown that total acellular ROS levels in e-cigarette aerosols are dependent on brand, flavor, operational voltage, and puffing protocol, but no studies so far have sought to investigate the role synthetic coolants have in modifying total acellular ROS levels in e-cigarette aerosols (Zhao, Zhang et al. 2018). In this study, we seek to understand the role WS-23 and WS-3 have in potentially modifying acellular ROS levels in e-cigarette-generated aerosols.

## 2. Materials & Methods

### 2.1. Procurement of e-liquid constituents and composition of e-liquid solutions

Propylene Glycol (PG), Vegetable Glycerin (VG), WS-23 solution (30% suspended in PG), and Koolada (10% WS-3 in PG) were purchased online from Flavor Jungle. 100 mg/mL nicotine salt solution (50:50 PG-to-VG ratio) was purchased online from PERFECTVAPE. E-liquid solutions comprising of PG, VG, salt nicotine, Koolada, and WS-23 were made. For our acellular ROS assays, the following e-liquids were made (Table 1).

**Table 1:** Composition of E-liquids Analyzed

| **Composition**<br>**of E-Liquid**<br>**Solution** | **PG:VG**<br>**Ratio (by**<br>**mass)** | **Nicotine**<br>**Concentration**<br>**(% by mass)** | **Cooling**<br>**Solution**<br>**Added** | **Cooling**<br>**Solution**<br>**Concentration**<br>**(% by mass)** |
| --- | --- | --- | --- | --- |
| PG:VG | 50:50 | 0.0 | None | 0.0 |
| PG:VG<br>(Nicotine) | 50:50 | 5.0 | None | 0.0 |
| PG:VG<br>+Koolada | 50:50 | 0.0 | FlavorJungle<br>Koolada (10%<br>WS-3 in PG) | 3.0 |
| PG:VG<br>+ WS-23 | 50:50 | 0.0 | FlavorJungle<br>WS-23 (30% in<br>PG) | 3.0 |
| PG:VG<br>(Nicotine)<br>+ Koolada | 50:50 | 5.0 | FlavorJungle<br>Koolada (10%<br>WS-3 in PG) | 3.0 |
| PG:VG<br>(Nicotine)<br>+ WS-23 | 50:50 | 5.0 | FlavorJungle<br>WS-23 (30% in<br>PG) | 3.0 |

### 2.2. Generation of Aerosols, Fluorescence Spectroscopy, and Acellular ROS Quantification

Each e-liquid solution was added to a new, empty refillable JUUL Pod (OVNStech, Shenzen, GD, China) (Mo: WO1 JUUL Pods) and aerosolized using a JUUL device (JUUL Labs Inc., Washington, DC, USA) (Mo: Rechargeable JUUL Device w/USB charger). Specifically, each JUUL device was attached to a Buxco Individual Cigarette Puff Generator (Data Sciences International (DSI), St. Paul, MN, USA) (Cat#601-2055-001), and subsequently, its component e-liquid was aerosolized and “bubbled” through 10mL of freshly made fluorogenic dye within a 50mL conical tube (Fig.1).

**Figure 1.**
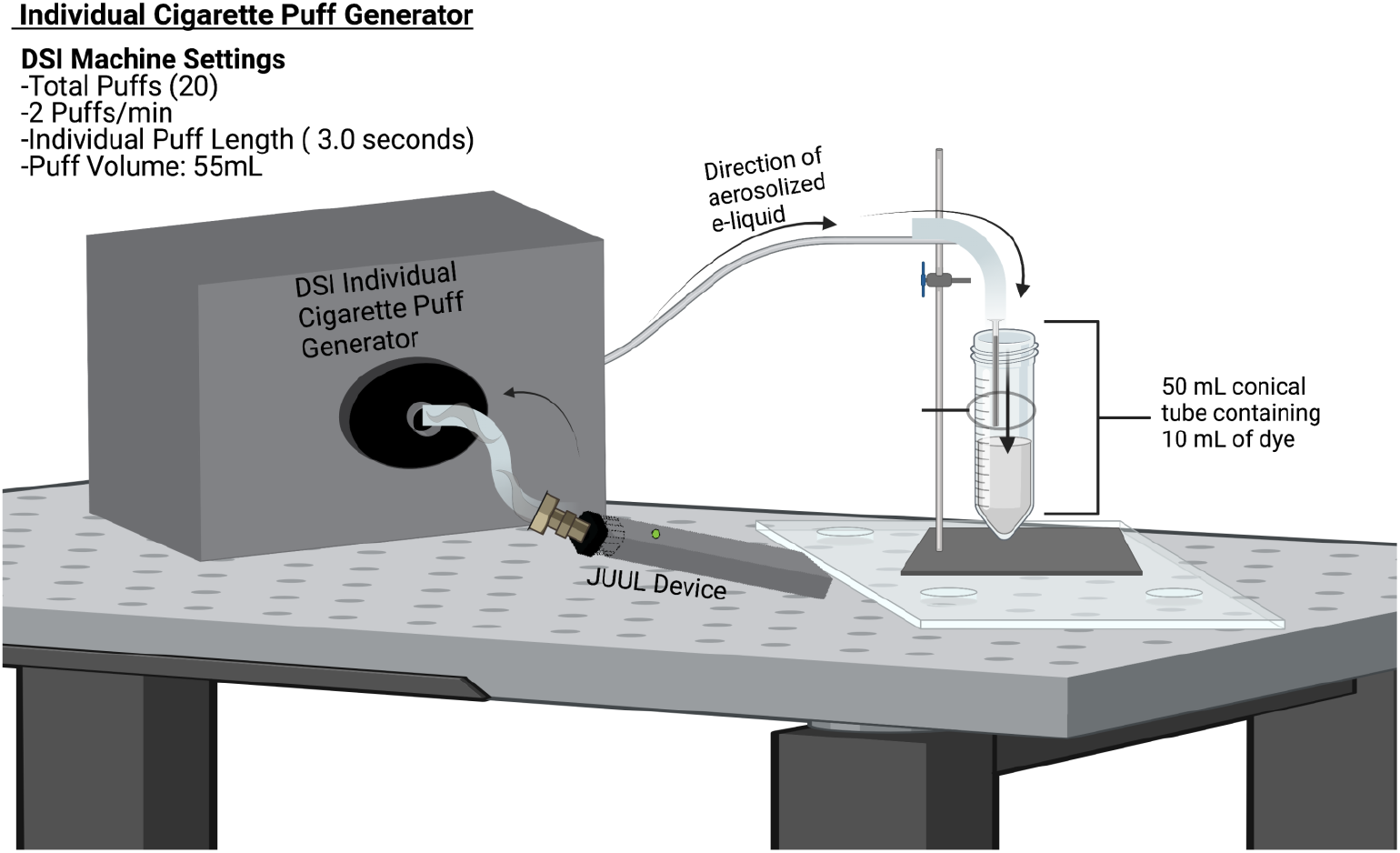
This pictogram shows the e-cigarette exposure generation system used in the study. E-cigarette aerosol was generated from the e-cigarette device using the artificial lung present in the Individual Cigarette Puff Generator. The e-cigarette aerosol then traveled to and was exposed to 10 mL of fluorogenic dye for one puff regimen at 1.5 L/min. One puff regimen consists of 20 total puffs (2 puffs/min) for 10 minutes, with the volume of each puff being 55.0 mL and each individual puff length lasting 3.0 seconds. Each conical tube was wrapped in aluminum foil to protect the fluorogenic dye from light. The entirety of the aerosolization and exposure process using the DSI machine was performed inside a chemical fume hood.

Cell permeant 2’,7’-dichlorodihydrofluorescein diacetate (H_2_DCFDA) (EMD Biosciences, San Diego, CA, USA) (Cat # 287810) dissolved in 0.01N NaOH, phosphate buffer, PO_4_, and horseradish peroxidase (Thermo Fisher Scientific, Waltham, MA, USA (Cat# 31491) were used to make the fluorogenic dye. The aerosols generated from each e-liquid solution were individually bubbled through 10 mL of H_2_DCFDA solution at 1.5 L/min. A schematic of the e-cigarette aerosolization procedure is shown in Figure 1. Each JUUL-pod containing a respective e-liquid solution had undergone three separate puffing regimens to create three separate samples of bubbled dye solution. The same puffing regimen was used for “bubbling” filtered air through fluorogenic dye for a negative control. For our positive control, the smoke generated from a research cigarette (Kentucky Tobacco Research & Development Center in the University of Kentucky, Lexington, KY, USA) (Mo: 3R4F) was bubbled through the fluorogenic dye. After “bubbling,” each resulting fluorogenic dye sample was placed in a 37 °C degree water bath (VWR 1228 Digital Water Bath) for fifteen minutes; subsequently, the solution was analyzed via fluorescence spectroscopy using a spectrofluorometer (Turner Quantech fluorometer, Mo. FM109535) in fluorescence intensity units (FIU). Readings on the spectrofluorometer were measured as H_2_O_2_ equivalents using a standard curve generated using the 0-50 µM H_2_O_2_ standards made.

### 2.3. Statistical Analyses

One-way ANOVA, unpaired t-test, and Tukey’s post-hoc tests were used for pairwise comparisons via GraphPad Prism Software version 8.1.1. Sample size was three. The results are shown as mean ± SEM. Data were considered to be statistically significant for *p* values < 0.05.

## 3. Results

### 3.1. Aerosolized nicotine-containing e-liquid with WS-23 contains significant levels of Acellular ROS

The levels of acellular ROS generated by the PG:VG solution (2.02-2.60 μM H_2_O_2_) were significantly higher than those generated by the filtered air control (0.96-1.66 μM H_2_O_2_) (Fig.2a). When the levels of acellular ROS generated by the PG:VG solution containing nicotine (5%) (1.13-1.84 H_2_O_2_ μM H_2_O_2_) and the filtered air control (0.96-1.66 μM H_2_O_2_) were compared, the generated ROS levels did not significantly differ (Fig.2b). The levels of ROS generated by the PG:VG with WS-23 solution (1.21-4.16 μM H_2_O_2_) did not significantly differ from those generated by the aerosolized PG:VG solution nor from the levels of acellular ROS generated by the filtered air control (Fig.3a). However, the levels of acellular ROS generated by the aerosolized e-liquid solution containing PG:VG with nicotine (5%) and WS-23 (3%) (1.94-2.95 μM H_2_O_2_) were significantly higher than those generated by the filtered air control (0.96-1.66 μM H_2_O_2_) (Fig.3b). In contrast, the levels of acellular ROS generated by the PG:VG solution containing nicotine and WS-23 (1.94-2.95 μM H_2_O_2_) did not differ significantly from those generated by the PG:VG solution containing nicotine (Fig.3b). When the levels of acellular ROS generated by the PG:VG solution containing nicotine and Koolada (2.27-2.57 μM H_2_O_2_) and the filtered air control were compared, the generated ROS levels were significantly different (Fig.4a). However, the difference in acellular ROS levels between aerosolized PG:VG with Koolada solution and aerosolized PG:VG solution was not significant (Fig.4a). Additionally, the levels of ROS generated by the PG:VG solution with nicotine and Koolada (1.79-3.35 μM H_2_O_2_) did not significantly differ from those generated by the aerosolized PG:VG with nicotine solution nor the filtered air control (Fig.4b).

**Figure 2.**
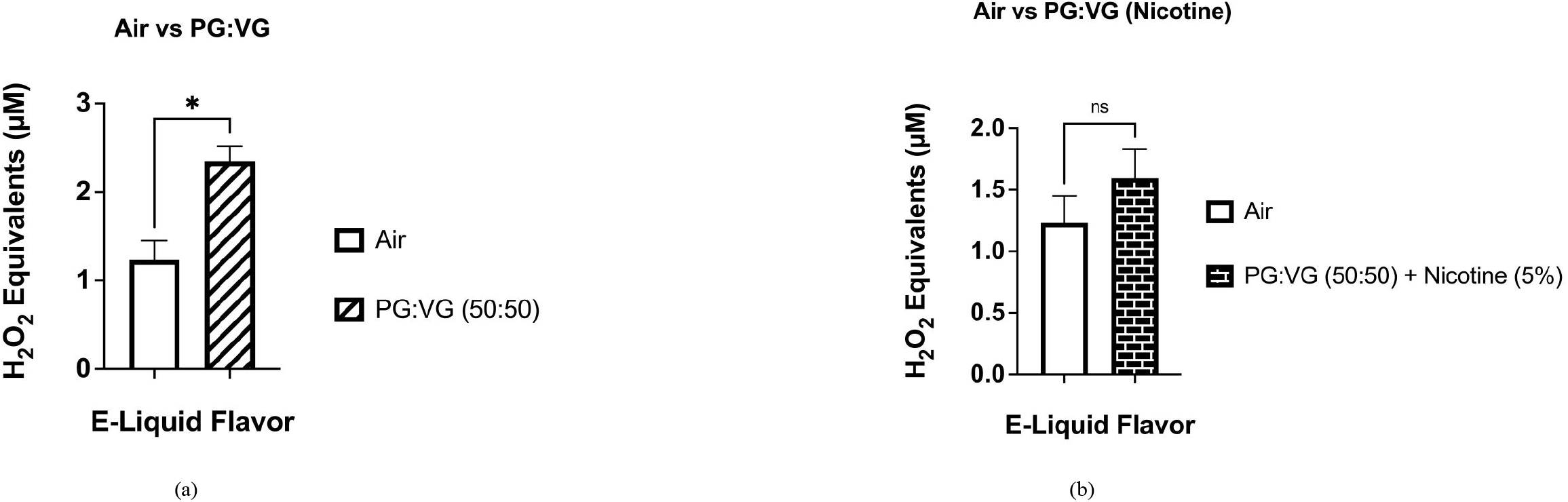
Comparisons between acellular ROS levels generated by aerosolized PG:VG (50:50), PG:VG (50:50) with nicotine, and a filtered air control. Acellular ROS was measured through hydrogen peroxide standards within aerosols generated from the previously mentioned e-liquids. Specifically, the e-liquid solutions were aerosolized using a JUUL device inserted into the Buxco Individual Cigarette Puff Generator. Data are represented as mean ± SEM, and significance was determined using an unpaired t-test. The ratio of PG:VG used in each solution and the percentage of nicotine each solution is made up of is listed above in the graphs. Smoke generated from a 3R4F research cigarette was used as a positive control.* p < 0.05 and ns is abbreviated for “Non-Significant” versus air control (*p* > 0.05). N=3

**Figure 3.**
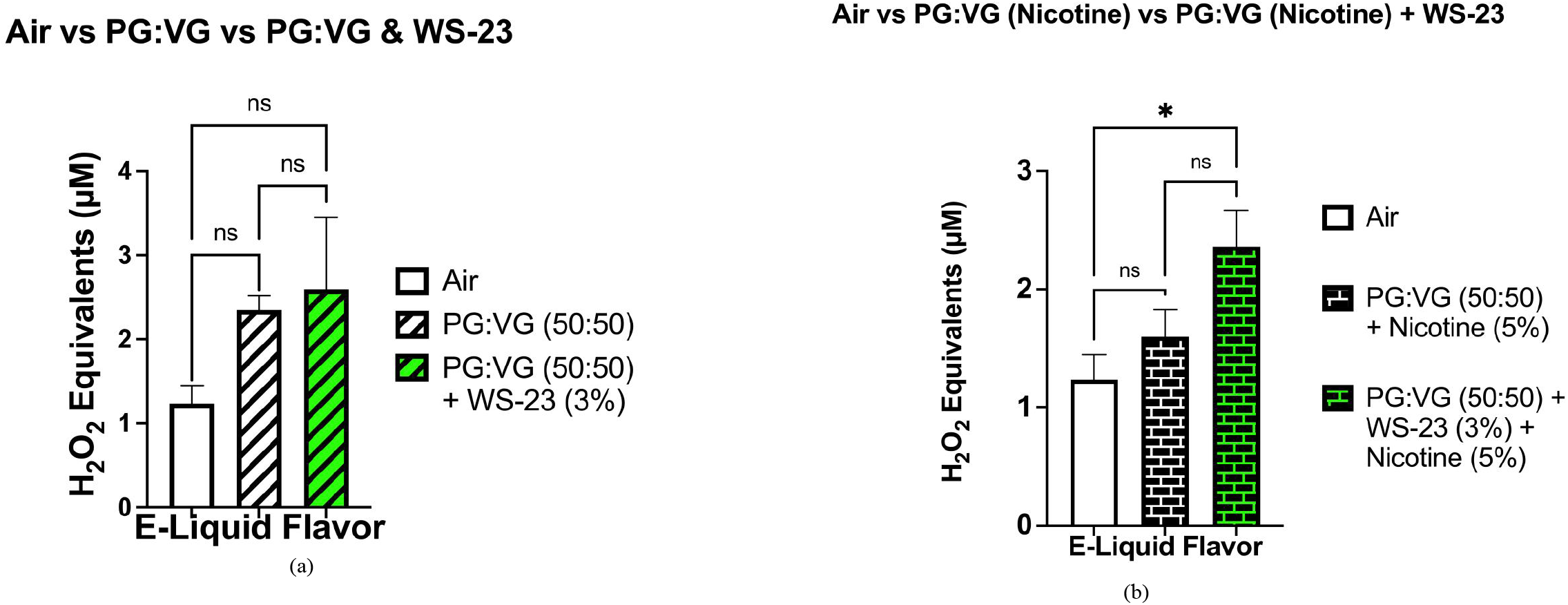
Comparisons between acellular ROS levels generated by aerosolized PG:VG (50:50), PG:VG (50:50) with nicotine, PG:VG (50:50) + WS-23, PG:VG (50:50) with nicotine + WS-23, and a filtered air control. Acellular ROS was measured through hydrogen peroxide standards within aerosols generated from the previously mentioned e-liquids. Specifically, the e-liquid solutions were aerosolized using a JUUL device inserted into the Buxco Individual Cigarette Puff Generator. Data are represented as mean ± SEM, and significance was determined using an unpaired t-test. The ratio of PG:VG used in each solution and the percentage of nicotine and WS-23 each solution is made up of is listed above in the graphs. Smoke generated from a 3R4F research cigarette was used as a positive control.* p < 0.05 and ns is abbreviated for “Non-Significant” versus air control (*p* > 0.05). N=3

**Figure 4.**
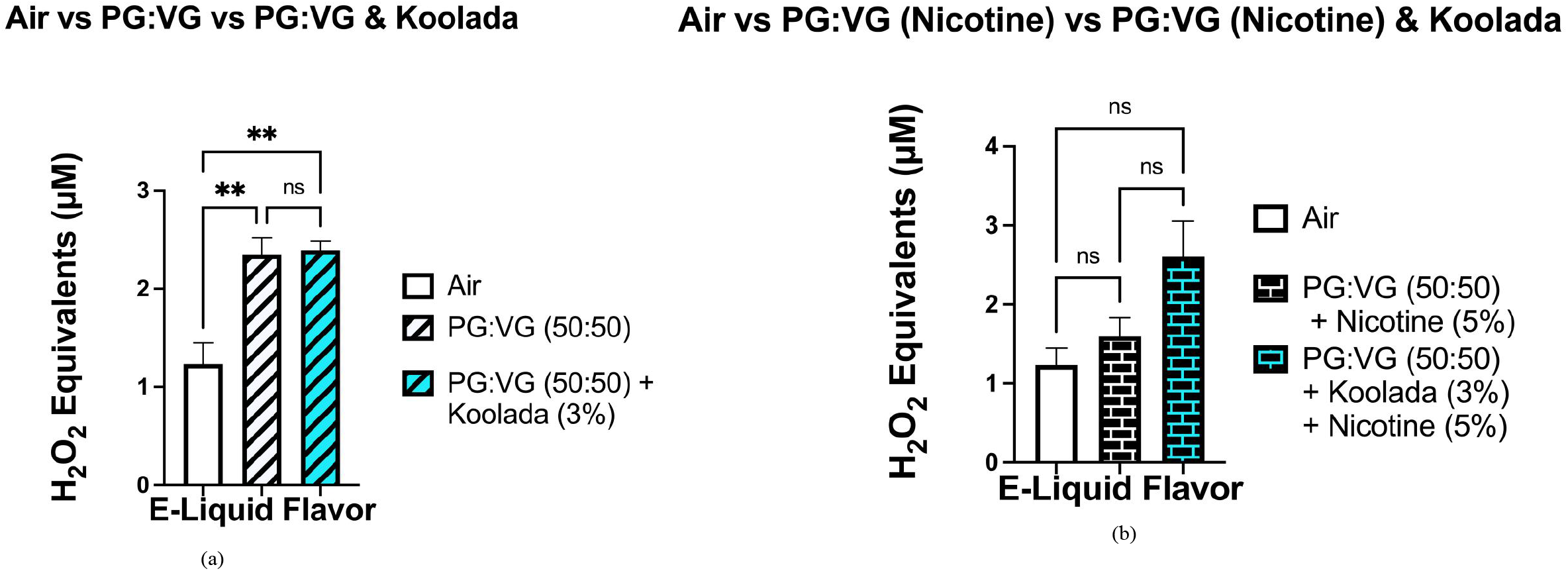
Comparisons between acellular ROS levels generated by aerosolized PG:VG (50:50), PG:VG (50:50) with nicotine, PG:VG (50:50) + Koolada, PG:VG (50:50) with nicotine + Koolada, and a filtered air control. Acellular ROS was measured through hydrogen peroxide standards within aerosols generated from the previously mentioned e-liquids. Specifically, the e-liquid solutions were aerosolized using a JUUL device inserted into the Buxco Individual Cigarette Puff Generator. Data are represented as mean ± SEM, and significance was determined using an unpaired t-test. The ratio of PG:VG used in each solution and the percentage of nicotine and Koolada each solution is made up of is listed above in the graphs. Smoke generated from a 3R4F research cigarette was used as a positive control.**p < 0.05 and ns is abbreviated for “Non-Significant” versus air control (*p* > 0.05). N=3

### 3.2 Koolada and WS-23 modify e-cigarette generated Acellular ROS Levels Similarly

The levels of acellular ROS generated by the PG:VG (50:50) with Koolada (3%) solution did not significantly differ from those generated by the PG:VG (50:50) with WS-23 (3%) solution (Fig 5.a). Additionally, neither the difference in acellular ROS levels between the aerosolized PG:VG with Koolada solution and the filtered air control nor between the aerosolized PG:VG with WS-23 solution and the filtered air control were significant (Fig.5a). When comparing the levels of ROS generated by the PG:VG with Koolada and nicotine solution to those generated by the PG:VG with WS-23 and nicotine solution, we see that they did not significantly differ (Fig.5b). Moreover, neither the difference in acellular ROS levels between aerosolized PG:VG with Koolada solution and the filtered air control nor between the aerosolized PG:VG with WS-23 solution and filtered air control were significant (Fig.5b). Our data shows that regardless of nicotine content (0% or 5%), minimal differences in acellular ROS levels exist when comparing the addition of Koolada and WS-23 to e-liquid base (PG:VG) (Fig.5a-b).

**Figure 5.**
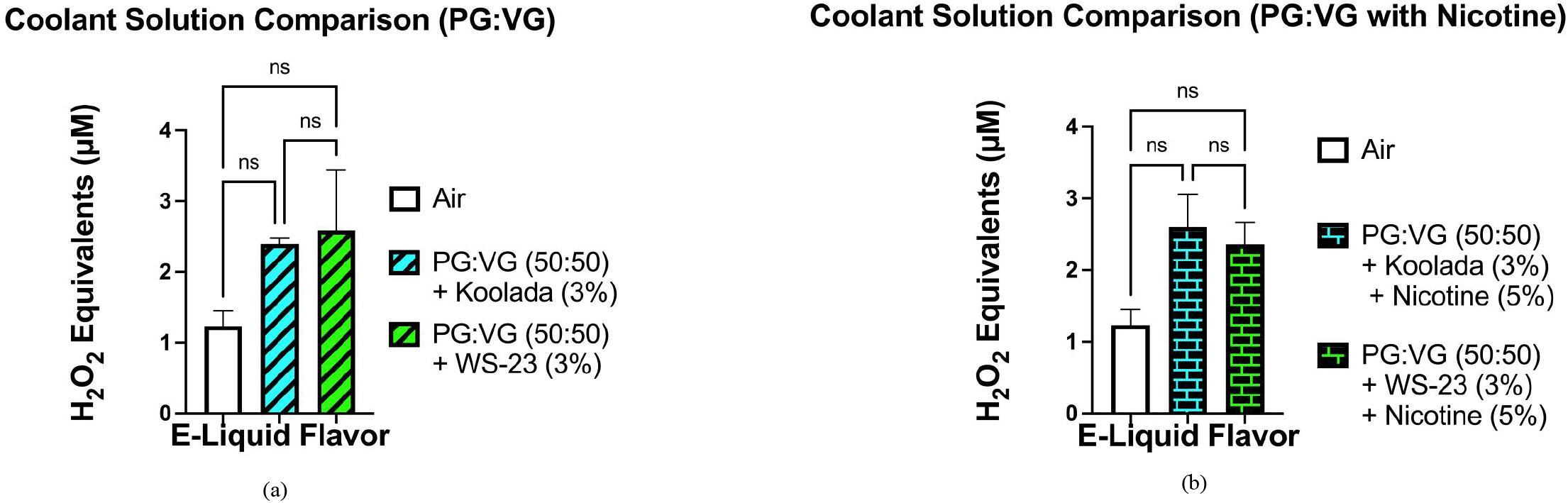
Comparisons between acellular ROS levels generated by aerosolized PG:VG (50:50) + Koolada, PG:VG (50:50) with nicotine + Koolada, PG:VG (50:50) + WS-23, PG:VG (50:50) with nicotine + WS-23, and a filtered air control. Acellular ROS was measured through hydrogen peroxide standards within aerosols generated from the previously mentioned e-liquids. Specifically, the e-liquid solutions were aerosolized using a JUUL device inserted into the Buxco Individual Cigarette Puff Generator. Data are represented as mean ± SEM, and significance was determined using an unpaired t-test. The ratio of PG:VG used in each solution, the percentage of nicotine, WS-23, and Koolada each solution is made up of is listed above in the graphs. Smoke generated from a 3R4F research cigarette was used as a positive control. ns is abbreviated for “Non-Significant” versus air control (*p* > 0.05). N=3

## 4. Discussion

With the surge of e-cigarette use amongst youth in the US in 2021 and the recent influx of “cool/iced” e-cig flavors in US marketplaces, there is a greater need to fill the knowledge gap on the safety of inhaling synthetic-coolant additives (Chen-Sankey, Bover Manderski et al. 2022). Our study sought to determine whether adding widely used synthetic coolants, WS-3 and WS-23, in e-liquids modifies the level of acellular ROS generated in e-cigarette aerosols. Our data suggest that the addition of WS-3 to e-liquid base (PG:VG), regardless of whether it contains 0% nicotine or 5.0% nicotine, has a minimal impact on modifying e-cigarette-generated acellular ROS levels. More specifically, neither the difference in acellular ROS levels between PG:VG with Koolada solution and PG:VG solution nor between PG:VG with Koolada and nicotine solution and PG:VG with nicotine solution were significant. Additionally, our data suggest that the addition of WS-23 to e-liquid base (PG:VG) with 5% nicotine does significantly impact e-cigarette-generated acellular ROS levels. To explain, the difference in generated acellular ROS levels between PG:VG with nicotine and WS-23 solution and the filtered air control was significant while that between the PG:VG with nicotine solution and filtered air control was not. Our data seems to suggest that synthetic coolants themselves have a limited impact in altering e-cig-generated acellular ROS levels generated from non-nicotine-containing e-liquids.

However, our findings concur with previous studies showing that aerosolized e-liquids contain significant levels of acellular ROS (Zhao, Zhang et al. 2018, Yogeswaran, Muthumalage et al. 2021). Regarding previous studies that analyzed acellular ROS levels within “cool/iced” flavored e-cigarettes, one study found differences in generated-acellular ROS levels between Tobacco-Derived Nicotine (TDN) and Tobacco-Free Nicotine (TFN) among cool/iced flavored e-cigarettes were minimal compared to tobacco and fruit flavors (Yogeswaran and Rahman 2022). In rodent studies, rats exposed to aerosolized e-liquid containing WS-23 at tested doses (via acute and subacute exposures) found no substantial changes in histopathologic analyses of vital organs nor relative organ weights (Wu, Liu et al. 2021). This same study, via a bronchioalveolar lavage fluid (BALF) analysis, found no significant difference in neutrophil concentration between rats which had undergone repeated 28-day WS-23 exposure and those apart of the respective control group (Wu, Liu et al. 2021). Neutrophils are major sources of endogenous ROS production.

Future studies aimed at understanding the role of WS-23 in modulating e-cig-induced oxidative stress should involve measurements of intracellular and extracellular ROS using isolated Polymorphonuclear Neutrophils (PMNs) (Kuhns, Priel et al. 2015). More specifically, PMNs isolated from blood collected from mice exposed to aerosolized e-liquids of varying WS-23 concentrations can be analyzed via luminol enhanced chemiluminescence exposure (Kuhns, Priel et al. 2015). The proposed experiment can provide insight into the differences between intra-and extra-cellular ROS of PMNs isolated from mice exposed to various concentrations of WS-23 (Kuhns, Priel et al. 2015). Regarding our understanding of the effects of other e-liquid coolant additives, using human bronchial epithelial cell cultures, one study found that treatment with menthol significantly increased mitochondrial ROS via the TRPM8 receptor (Nair, Tran et al. 2020).

Regarding limitations in our study, our study did not include the treatment of airway epithelial cells with aerosolized e-liquids. Previous studies have shown that treatments with e-liquids induce significant levels of ROS production in Human Bronchial Epithelial cells (BEAS-2B) (Wang, Wang et al. 2021). Epithelial cells lining the airways are the first structural cell targets of any inhaled substances (Hiemstra, Tetley et al. 2019). Likewise, a better understanding of how synthetic coolants modulate e-cigarette-induced oxidative stress in the lungs can be obtained through cellular ROS assays. More specifically, future studies should conduct a staining MitoSress assay using airway epithelial cells exposed to aerosolized e-liquids containing various concentrations of synthetic coolants (WS-3 and WS-23) (Muthumalage, Lamb et al. 2019). Through this proposed assay, an understanding of how exposure to aerosolized synthetic coolants affects mitochondrial ROS production can be obtained. However, our study has shown that the addition of WS-3 and WS-23 to e-liquids has a minimal effect on modifying acellular ROS levels within aerosolized non-nicotine-containing e-liquid base. Thus, these preliminary findings indicate the need for further evaluation on the potential health risks associated with inhaling newly marketed e-cigarettes containing synthetic coolants. Specifically, our findings highlight the need for further investigation into the role of WS-3 and WS-23 in disrupting the endogenous oxidant and antioxidant balance in airways upon inhalation.

## Acknowledgments

Figure 1 and the Graphical Abstract were made using BioRender and AdobeIllustrator. Figures 2,3,4, and 5 were made using GraphPadPrism. Joseph Lucas (JL) and Dr. Thivanka Muthumalage (TM) for technical help and discussions.

## Author Contributions

Conceptualization, I.R.; methodology, I.R.; assay performance: S.Y, software, S.Y.; validation, S.Y, I.R.; formal analysis, S.Y.; investigation, S.Y.; re-sources, I.R.; data curation, S.Y.; writing—original draft preparation, S.Y and I.R.; writing—review and editing, S.Y., M. M., H.S.C., and I.R.; visualization, S.Y.; supervision, I.R.; project administration, I.R.; funding acquisition, I.R. All authors have read and agreed to the published version of the manuscript.

## Data Availability Statement

We declare that we have provided all the data in figures.

